# In silico design of ligand triggered RNA switches

**DOI:** 10.1101/245464

**Authors:** Sven Findeiß, Stefan Hammer, Michael T. Wolfinger, Felix Kühnl, Christoph Flamm, Ivo L. Hofacker

## Abstract

This contribution sketches a work flow to design an RNA switch that is able to adapt two structural conformations in a ligand-dependent way. A well characterized RNA aptamer, i. e., knowing its *K*_*d*_ and adaptive structural features, is an essential ingredient of the described design process. We exemplify the principles using the well-known theophylline aptamer throughout this work. The aptamer in its ligand-binding competent structure represents one structural conformation of the switch while an alternative fold that disrupts the binding-competent structure forms the other conformation. To keep it simple we do not incorporate any regulatory mechanism to control transcription or translation. We elucidate a commonly used design process by explicitly dissecting and explaining the necessary steps in detail. We developed a novel objective function which specifies the mechanistics of this simple, ligand-triggered riboswitch and describe an extensive *in silico* analysis pipeline to evaluate important kinetic properties of the designed sequences. This protocol and the developed software can be easily extended or adapted to fit novel design scenarios and thus can serve as a template for future needs.

## 1. Introduction

Riboswitches are highly structured RNA sequences commonly found in the 5’-untranslated region (UTR) of prokaryotic messenger RNAs (mRNAs). Within this regulatory domain, they are responsible for altering gene expression on the transcriptional or translational level in response to environmental changes, which is typically the concentration of a small ligand [1]. A riboswitch consists of two components: i) a sensory domain and ii) a regulatory domain. While the former specifically senses the environmental change, the latter is responsible for influencing the expression level of the downstream gene. Beside those switches that follow this commonly assumed two-component model, examples are known where a sensory domain alone, i. e., an aptamer, is able to alter gene expression [2, 3, 4, 5]. The possibility to encode effective sensors at RNA level makes riboswitches valuable gadgets that can directly interfere with the complex process of gene expression without the need of additional co-factors such as proteins. Here we elucidate the complex process of designing such a ligand-sensing riboswitch that, for simplicity, does not implement a specific regulation mechanism at transcriptional or translational level. The RNA sequence should “simply” adapt two alternative conformations depending on the presence or absence of a ligand. Therefore, we aim to extend an aptamer such that an alternative structural conformation is formed in the absence of the ligand.

Successful design approaches show that the problem of generating an artificial RNA sequence exhibiting a prescribed functionality needs to be formulated as a multi-step approach, including computational and experimental, analytic and constructive methods [6, 7]. Early design publications already followed such a multi-step scheme but included manual steps instead of computational methods, as there were just no computational tools available that implemented the features actually needed by the experimentalists. However, this changed over time and recently the common trend is to perform as many steps as possible with the support of advanced *in silico* methods [8].

As a first step, it is important to analyze the underlying biological system, the cellular environment and, most importantly, all the properties of the building blocks to use. With this information, it is then possible to design a model describing the functionality of the novel RNA molecule in its environment. To determine a sequence with the requested characteristics, usually an optimization problem is formalized, where the objectives are specified as constraints and a mathematical function describing various biophysical properties of the system. Obtaining a sequence compatible with constraints such as specific target structures and sequence motifs is a quite tricky task, which was solved with various methods ranging from manual design [9] to graph-theoretical coloring algorithms as implemented in *RNAblueprint* [10]. A variety of well-established optimization methods such as the Metropolis – Hastings algorithm [11, 12] or genetic algorithm based approaches [13] were used to find optimal solutions by traversing through the constrained solution space [14]. However, until now, only little effort was made to find proper objective functions. Often only some static properties of the molecule’s energy landscape are used instead of more directly characterizing the mechanism of the artificial device. Only recently, some published design programs started to allow to compile an objective function from a catalog of predefined functions [15, 13]. *RNAblueprint* [10] went one step further and allowed to formulate the objective utilizing a scripting interface, which gives the user complete control over the optimization procedure.

To narrow down the number of obtained RNA sequences, a subsequent step to analyze and filter the obtained solutions was almost always performed. Essentially, the differences and advantages of various solutions are explored. The generation of proper visualizations or the evaluation of additional properties that could not be incorporated into the objective function help to perform this selection process.

Finally, it is crucial to biologically test for the desired functionality of the designed molecule as many biologically relevant aspects cannot be easily included in the objective of the optimization approach.

In this contribution, we aim to closely follow the described design steps to generate a simple, ligand-triggered riboswitch, see Figure 1. Therefore, we combine the previously published RNA design software *RNAblueprint* with analysis tools like the coarse graining program *barriers* and the kinetics simulator *treekin*, see Table 1. To achieve our goal, we propose a functional model, specify valid constraints, and develop an objective function, which directly describes the functionality of this riboswitch at an abstract level. To compute the quality measures used by our objective, we resort to the well-established thermodynamic RNA folding model implemented in the *ViennaRNA* package. An extensive *in silico* analysis pipeline evaluates important properties of the designed sequences and thus helps to narrow down and filter the list of obtained sequences that might be sent to the laboratory for biological testing. Although we describe a work flow for the purpose of generating a *specific* riboswitch, the overall result comprises the developed protocol and software, which can easily be extended or adapted to fit novel design scenarios and thus could serve as a design strategy scaffold.

**Table 1:**
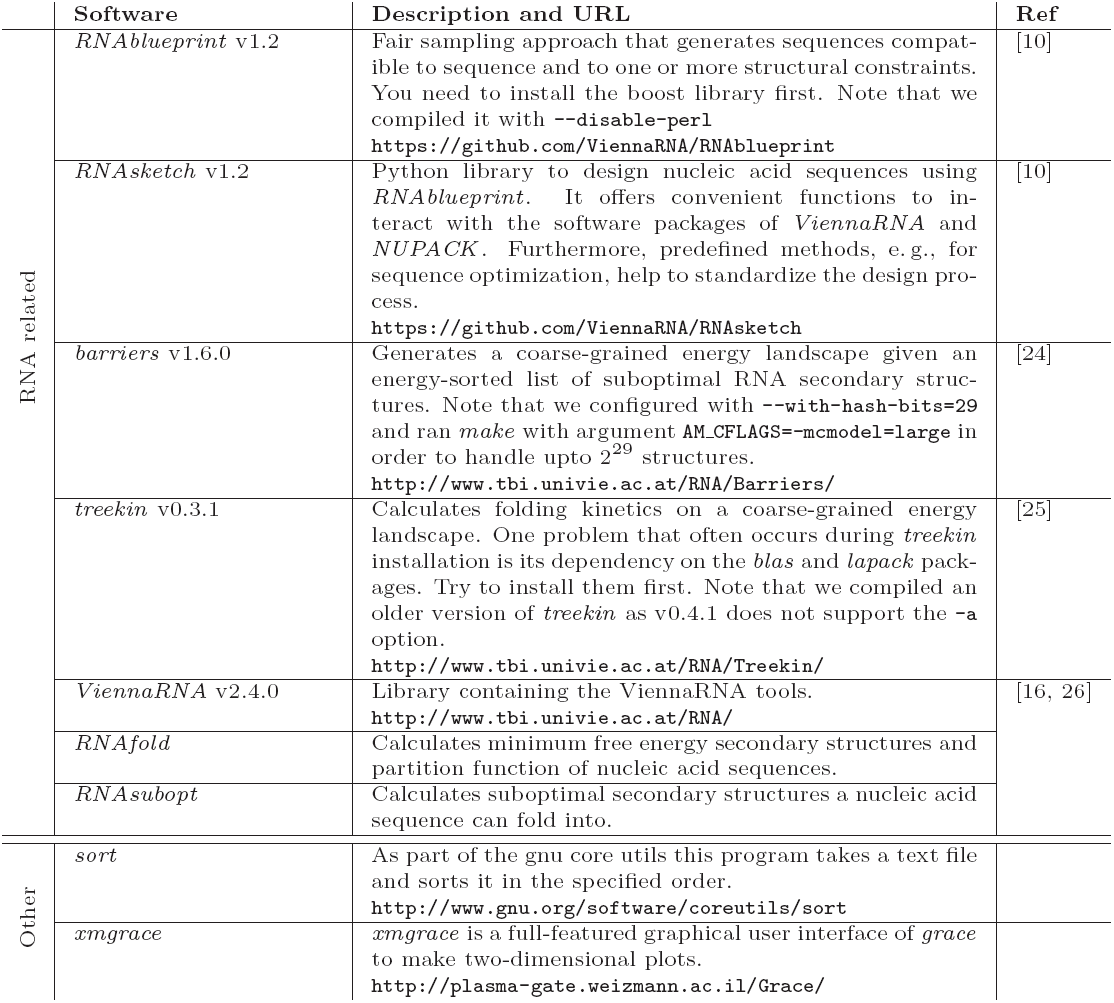
Summary of the utilized software. RNA related software tools are either standalone or part of the *ViennaRNA* package. The installation procedure is documented on the web pages listed. Standard Unix tools are tagged as “Other” and are typically included in or easy to install with the package manager of any distribution.

**Figure 1:**
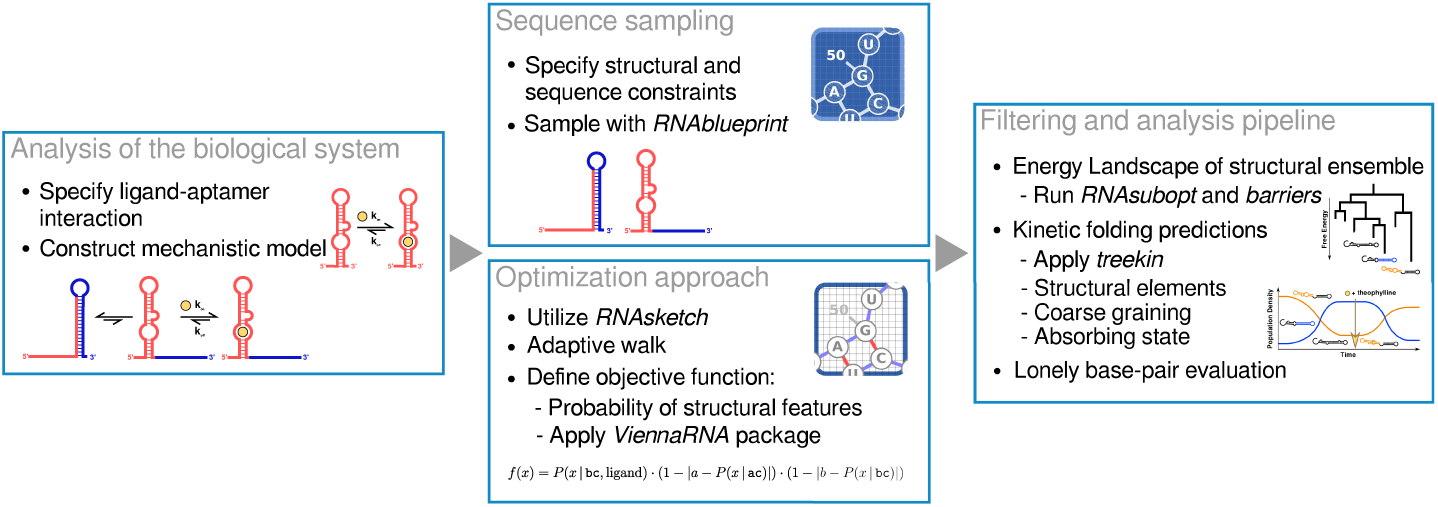
Graphical summary of the applied design approach. The work flow is depicted from left to right and consists of three major steps. For the individual parts important keywords and utilized software is listed and we refer to the individual sections of this contribution for more detailed information.

## 2. Materials and Methods

### 2.1. Specifying the design constraints

Given a model describing a desired riboswitch or functional RNA, it now needs to be converted into a machine-readable format in order to computationally generate valid sequences. Thus, the desired properties and the functionality can be expressed as a combination of constraints such as structural requirements and various properties specifying the energy landscape, and the kinetic folding properties. We specified these constraints of the functional states in the file design_input.txt:

~~~
# alternative conformation:
…………………….(((((((((((……)))))))))))………..
# binding competent conformation: (((((…((((((((…..)))))…)))…)))))..………………….
# sequence constraint:
AAGUGAUACCAGCAUCGUCUUGAUGCCCUUGGCAGCACUUCA NNNNNNNNNNNNNNNNNNNNNN
~~~

These sequence and structure constraints are represented in IUPACK and dot – bracket notation, respectively. In the sequence constraint, A, U, G and C correspond to the nucleotides adenine, uracil, guanine and cytosine, respectively. To positions marked with N, any nucleotide can be assigned as long as they are compatible to the structural constraints, where “.” represents an unconstrained position and matching brackets “()” two positions paired with each other. Please note, constraints resulting from or overlapping with the chosen aptamer are separated by a space that needs to be removed when constraints are used as input for *RNAblueprint*, the utilized sequence sampler. We collected all applied tools in Table 1 including a short summary, the download link, citation and further remarks. *RNAblueprint* can be invoked by executing the following command:

~~~
$ RNAblueprint-v < design_input.txt > design_output.txt
~~~

This returns how many compatible sequences exist (1.34218 × 10^8^ for the given example) and, by default, ten randomly generated sequences, which are written into file design output.txt. In principle, each of these sequences can fold into both specified structures, the most stable structure typically being a hybrid. Please note, that the obtained sequences are randomly generated and thus vary on every call.

For the demonstration of our analysis workflow, we selected a sequence that exhibits interesting properties, although it was not the best design generated during the applied optimization procedure. During such an optimization run thousands of compatible sequences are evaluated with respect to an objective function.

### 2.2. Prediction of minimum free energy structures

A transcribed RNA molecule immediately forms intra-molecular base pairs, folding into a structural conformation known as its (secondary) structure. The structure, in turn, often determines the RNA’s biological function, e. g., in our case, the binding affinity for a given ligand. Any structure of a given RNA sequence can be assigned an energy value — the Gibbs free energy — and the structure expressed most likely is the one having the lowest possible energy. It is therefore called the minimum free energy (MFE) structure.

To predict the MFE of a given sequence and its associated secondary structure, we use the tool *RNAfold* included in the *ViennaRNA* package (cf. Table 1). We first store an example sequence in a text file exa.txt and subsequently apply *RNAfold* to it.

~~~
$ echo “AAGUGAUACCAGCAUCGUCUUGAUGCCCUUGGCAGCACUUCAGUUGCUGGGGGAAUGUUUUUGU” \
> exa.txt
$ cat exa.txt | RNAfold
AAGUGAUACCAGCAUCGUCUUGAUGCCCUUGGCAGCACUUCAGUUGCUGGGGGAAUGUUUUUGU
………..(((((…..)))))((((((((((……))))))))))………… (−19.70)
~~~

The above invocation of *RNAfold* returns, beside the input sequence, its most stable structure in dot – bracket notation and the corresponding MFE. Energies are given in kcal mol^−1^.

### 2.3. Modeling ligand binding with soft constraints

To incorporate the ability of binding a specific ligand into an *in silico* RNA design process, a model aware of the stabilizing contributions of this dimerization on the resulting RNA – ligand complex is required. For the work presented here, the recently implemented soft constraint framework of the *ViennaRNA* package [16] has been applied. Among other things, it allows to add an energy bonus to structural states that exhibit a certain motif. This enables for a direct integration of the effects of ligand binging into the RNA structure prediction process [16]. When evaluating the structure ensemble of a given molecule containing the theophylline aptamer sequence, an energy bonus of Δ*G* = −9.22 kcal mol^−1^ is added to every secondary structure that contains the correctly folded binding pocket. This value is obtained from the relation Δ*G* = *R* × *T* × ln *K*_*d*_ for the gas constant *R* = 1.987 17 cal mol^−1^, the temperature *T* = 310.15 K, and the experimentally measured dissociation constant *K*_*d*_ = 0.32 μM [17]. Using the example sequence and the --motif option of *RNAfold*,

~~~
$ cat exa.txt | RNAfold-p \
--motif = “GAUACCAG&CCCUUGGCAGC,(…((((&)…)))…),-9.22”
AAGUGAUACCAGCAUCGUCUUGAUGCCCUUGGCAGCACUUCAGUUGCUGGGGGAAUGUUUUUGU
.((((…((((((((…..)))))…)))…))))((((…))))………….. (−21.92)
,((((…((((((((,…,)))))…)))…))))|(((…})),………….. [−23.32]
.((((…((((((((…..)))))…)))…)))).(((…)))…………… {-12.20 d = 4.04}
frequency of mfe structure in ensemble 0.103202; ensemble diversity 6.30
~~~

the MFE structure now contains the binding-competent aptamer fold with a corrected energy value (cf. first row after the sequence). As a result from using the-p option, a condensed representation of the base pair probabilities of each nucleotide in the ensemble with the Gibbs free energy of the soft-constrained ensemble *G*(*x* | *s*) (second row) as well as the centroid structure, i. e., the consensus structure of all base pairs with a probability higher than 50% in the ensemble [18], and its free energy (third row) are printed.

### 2.4. Obtain the probability of structural features

In a design process, it is usually desired to enforce the presence or absence of certain structural motifs, or even requires a certain sub-structure to be present with a specific probability. This requires a method that can determine the fraction of structures of a given RNA sequence that contain a given motif. An objective function can then use this information to compute probabilities of motifs and accordingly select sequences suitable for the design goal.

*Hard constraints* of the *ViennaRNA* package are well suited for such tasks. They allow to restrict the conformations of an RNA to states containing a combination of unconstrained bases “.”, bases that have to be unpaired “x”, bases that have to be paired no matter to which binding partner “|” and base pairs indicated by matching brackets “()”. It is furthermore possible to specify if a base has to be paired with a binding partner up-or downstream by “<” and “>“, respectively. Note, that structures lacking some constraints are also counted as long as no base pair conflicts with the constraint. To only include structures possessing all specified base pairs, use the --enforceConstraint option. To calculate the probability of the alternative conformation, one can use the following constraint and command:

~~~
constraint.txt AAGUGAUACCAGCAUCGUCUUGAUGCCCUUGGCAGCACUUCAGUUGCUGGGGGAAUGUUUUUGU
…………………….(((((((((((……)))))))))))………..
$ cat constraint.txt | RNAfold-C-p--canonicalBPonly\
--motif = “GAUACCAG&CCCUUGGCAGC,(…((((&)…)))…),-9.22”
AAGUGAUACCAGCAUCGUCUUGAUGCCCUUGGCAGCACUUCAGUUGCUGGGGGAAUGUUUUUGU
…………((((…..))))(((((((((((……)))))))))))……….. (−19.50)
….,…….((((|…,))))(((((((((((……)))))))))))……….. [−20.19]
…………((((…..))))(((((((((((……)))))))))))……….. {-19.50 d = 3.89}
frequency of mfe structure in ensemble 0.32628; ensemble diversity 6.02
~~~

This performs a constrained (-C) partition function (-p) fold simulating lig-and binding (--motif). The --canonicalBPonly option removes non-canonical base pairs, e. g., U-U, from the structure constrain if they where erroneously added. Here, all structures of a sequence *x* containing only base pairs compatible to the hard constraint *h* and the soft constraint *s* are considered during the calculation. The Gibbs free energy *G*(*x* | *h, s*) of those structures can then be used to calculate the frequency of the constrained sub-structure within the complete ensemble, which is denoted as

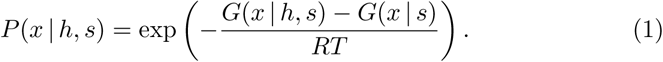

For more background information on the relation of these energies and the probabilities, please refer to subsection 2.6. Re-running the last command without the-C option yields the Gibbs free energy *G*(*x* | *s*) without the hard constraint *h*. This will include all suboptimal structures in the calculation, but still uses the soft constraint option to model ligand binding (cf. subsection 2.3). For the example above, *G*(*x* | *s*) and *G*(*x* | *h, s*) are −23.32 kcal mol^−1^ and −20.19 kcal mol^−1^, respectively, and the resulting probability is

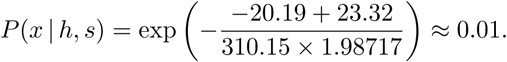

The frequency of a structural motif in the absence of any ligand can be obtained by running both commands without the --motif option. Thus, we can also calculate *P* (*x* | *h*) as denoted in Equation 1.

### 2.5. Enumerating suboptimal structures of an RNA molecule

To analyze the *kinetics* of an RNA molecule, at least a part of its structural ensemble needs to be explicitly constructed. This is a challenging task as the number of structures even for small RNAs is enormous. The tool *RNAsubopt* (cf. Table 1) can be applied to generate all structures a given sequence can adopt up to a given energy threshold. Consider the following example:

~~~
seq1.txt
AAGUGAUACCAGCAUCGUCUUGAUGCCCUUGGCAGCACUUCAGUUGUUGAGGGGGCUCAAUGAC
$ cat seq1.txt | RNAsubopt -e 1.2 –s
AAGUGAUACCAGCAUCGUCUUGAUGCCCUUGGCAGCACUUCAGUUGUUGAGGGGGCUCAAUGAC −21.60 1.20
…………….((((((((.(((((((((((……))))))))))).).)))).))) −21.60
….(((……)))((((((((.(((((((((((……))))))))))).).)))).))) −21.50
(((((…((((((((…..)))))…)))…)))))..(((((((((….))))))))) −21.10
(((((…(((((((((…))))))…)))…)))))..(((((((((….))))))))) −20.80
.((((…((((((((…..)))))…)))…))))…(((((((((….))))))))) −20.80
..(((…….))).((((((((.(((((((((((……))))))))))).).)))).))) −20.60
.((((…(((((((((…))))))…)))…))))…(((((((((….))))))))) −20.50
..((…….))…((((((((.(((((((((((……))))))))))).).)))).))) −20.50
…(((..((.(((((…..)))))((((((((((……))))))))))))..)))….. −20.40
~~~

where all suboptimal structures with an energy at most 1.2 kcal mol^−1^ above the MFE are generated. Note that the number of generated structures grows exponentially with both the sequence length and the size of the selected energy and. Thus, larger instances of these calculations do not only consume CPU time, but also generate files of several gigabytes in size. To reduce the number of generated sequences, it is possible to skip all structures containing *lonely base pairs*, i. e., helices of length one, by applying the --noLP option of *RNAsubopt*, cf. subsection 2.6.

As the sorting routine applied by *RNAsubopt* (-s option) might fail on huge instances even on high memory machines with giga-or even terabytes of RAM, a workaround is to pipe the *RNAsubopt* output to Unix’s *sort* tool. The latter scales much better with the memory consumption of the typically huge ensemble sizes. The following example generates the full *RNAsubopt* outputs. The execution of this command may take some time as the *RNAsubopt* output of the example sequence is approximately 16 GB in size.

~~~
$ cat seq1.txt | RNAsubopt-e 22.60 | sort-k2,2n-k1,1r-S20G > seq1.sub
~~~

Here, the main memory buffer allocated by *sort* is set to 20 GB. Above this threshold, *sort* will dump data to temporary files on the hard drive. Assuming enough disk space is available, this still implies performance loss but makes it possible to process even huge *RNAsubopt* output. We estimated the energy band width to use for the-e option by folding the sequence with *RNAfold* (cf. subsection 2.2), setting its value to −1 × MFE + 1 to convert the minimum free energy (MFE) into a positive value and also take a few structures with positive energies into account. The obtained file seq1.sub contains a list of all possible structures within 22.60 kcal mol^−1^, sorted by ascending energy values.

### 2.6. Assessing the impact of avoiding lonely pairs

The --noLP option of *RNAsubopt* (and *barriers*, cf. subsection 2.7) achieves a considerable speed-up by neglecting structures containing so-called *lonely base pairs*, i. e., base pairs which are not directly surrounded by another base pair. Put differently, this option enforces a minimal helix length of two base pairs. The biological motivation of this optimization is that lonely base pairs usually destabilize a secondary structure and thus would open up again quickly. Structures not containing lonely pairs are called *canonical* structures.

Using --noLP significantly reduces the resources required for conducting the analysis, but may also bias its results. Therefore, when analyzing a newly designed sequence, the question arises whether applying this heuristics will, in this specific case, yield accurate results or not. Here, we derive a measure that helps to answer this question for individual sequences.

The probabilities of the secondary structures for a given RNA sequence *x* follow a Boltzmann distribution, i. e., the probability of a secondary structure *φ* is proportional to 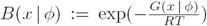. Here, *R* is the universal gas constant, *T* is the absolute temperature, and *G*(*x* | *φ*) is the Gibbs free energy of the RNA *x* folded into the structure *φ*. The term *B*(*x* | *φ*) is referred to as the *Boltzmann weight* of *φ*. If Φ is the entire structure ensemble of *x*, then the *partition function* of *x* is given by *Z* = ∑_*φ*∊φ_*B*(*x* | *φ*), and the probability of structure *φ* in the ensemble is

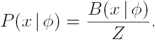

Note that 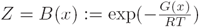, i. e., the partition function is the Boltzmann weight of the ensemble energy *G*(*x*).

It is reasonable to assume that leaving out extremely unlikely structures will not significantly change the results of the analysis to be performed, so one way to assess the impact of the heuristics is to enumerate structures up to a certain energy threshold and compute the fraction of structures that contain lonely pairs and will therefore be excluded from the simplified analysis. Furthermore, instead of simply taking the fraction of *counts* of structures with and without lonely pairs, one can get more profound results by comparing the *sums of their Boltzmann weights* corresponding to the probabilities of the respective sets of structures.

To achieve this for a given sequence *x*, first calculate the partition of the full ensemble *Z* = *B*(*x*) using the ensemble energy *G*(*x*) that can be computed by running RNAfold-p. Now, RNAsubopt-e can be used to enumerate structures within a given energy band above the MFE. Initialize variables *Z*^(0)^ ← 0 and 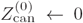, which will be used to store the approximations of the partition functions of the full and the canonical ensemble, respectively. For the *t*-th output structure *φ* of *RNAsubopt*, compute its Boltzmann weight *B*(*x* | *φ*) and set *Z*^(*t*)^ ← *Z*^*t*-1^ + *B*(*x* | *φ*). Then, verify whether *φ* is canonical and, if this is the case, set 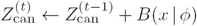, otherwise, leave it unchanged by setting 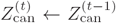. Finally, compute the fractions 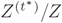 and 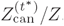, where *t*^*^ is the final value of *t*. The first fraction measures the structure coverage, i. e., which Boltzmann-weighted fraction of structures has been analyzed. It should be close to 1 for reliable results and can be improved by increasing the width of the energy band that limits the structure enumeration. The second fraction approximates the ratio of canonical structures in the ensemble.

As a side node, it is arguable that instead of the more complex enumeration process just described, the ensemble energy of the canonical ensemble could be directly computed using RNAfold-p --noPS. However, due to current technical limitations, the returned ensemble energy is only an upper bound of the actual value and may dramatically over-predict the fraction of canonical structures.

### 2.7. Analyzing the high-dimensional structure landscape

As the number of structures for a given RNA sequence grows exponentially with the sequence length, the folding process cannot be simulated with every single secondary structure even for small RNAs. Therefore, the number of simulation states needs to be reduced to a feasible number, ideally without biasing the outcome. This can be achieved by applying a coarse graining approach, which reduces the size of the high-dimensional structure landscape the sequence spans to a much smaller set of *macro-states*, each of which represents a set of multiple structures.

The tool *barriers* (cf. Table 1) implements a flooding algorithm that effectively coarse grains an energy landscape to macro-states or *basins*, each represented by a local minimum of the folding landscape. Each basin contains all the structures connected to its representative local minimum by the folding path of steepest descend. For each two macro-states, the tool also computes *barrier height*, i. e., the highest intermediate structure (with respect to its energy) that has to be overcome in order to refold from one state to the other. It can be used to visualize the RNA landscape by drawing a *barrier tree*.

As input *barriers* requires a list of all suboptimal structures within a certain energy range, sorted by ascending energy value. How to obtain such a list is explained in subsection 2.5. To obtain correct simulation results, the energy range has to be large enough to connect all generated macro-states. If this is not the case, the width of the energy band has to be increased. Alternatively, heuristic approaches such as *findPath* [19] may be used to connect formerly disconnected states. In order to handle the huge amount of structural states generated by our design example, it is mandatory to configure barriers using the option --with-hash-bits = 29 and to run *make* with the argument AM CFLAGS =-mcmodel = large.

Once the input file has been generated, *barriers* can be applied to it by executing

~~~
$ barriers --max = 500-G RNA-M noShift --bsize--rates < seq1.sub > seq1.bar
~~~

The --max = 500 option specifies the number of macro-states to be generated,-G, specifying the graph type, is set to RNA,-M noShift disallows so-called shift moves (i. e., a move changing exactly one of the two indices of an existing base pair), and --bsize and --rates enable the output of the size of each basin, and to compute transition rates between these macro-states. The results of *barriers* are then piped into the file seq1.bar. A graphical representation of the barrier tree in the PostScript format is by default saved to a file named tree.ps, whereas the rates are stored in file rates.out.

Note that *barriers* needs to be run with the -G RNA-noLP option when predicting an ensemble without lonely pairs.

### 2.8. Simulating kinetic folding using macro-states

When relying solely on thermodynamic criteria — e. g., probabilities of given conformations — during an RNA design process, one may miss important traits of the candidate sequences. Transcriptional riboswitches, for example, interact with the RNA polymerase in a time-critical manner, and information about the presence or absence of specific sub-structures within certain time frames are necessary to ensure correct switching behavior Such knowledge can be obtained by running a kinetics simulation for the given RNA sequence. As this type of analysis is too time-consuming to be included into the design process directly, it should be performed on a small set of promising candidates to verify their functionality.

The program *treekin* can be used to simulate single-molecule kinetics, which solves a continuous-time Markov process by numerical integration with the infinitesimal generator being a rate matrix. The latter is obtained by running *barriers*, which estimates the transition rates from each macro-state to all the other ones and stores them as a matrix in the file rates.out (subsection 2.7). The computation is performed using *treekin* as follows:

~~~
$ treekin --p0 1 = 1-m I-f rates.out --t8 = 1E12 < seq1.bar > seq1.tkin
~~~

Here-m I tells *treekin* to parse the file specified by -f as *barriers* output, --t8 sets the maximum simulation time to 1 × 10^12^ arbitrary time units (AU) and --p0 sets the initial population size of the selected minimum of the barrier tree. Here, we set the global minimum of the barrier tree (i. e., macro-state 1) to be 100%. The output can then be visualized by using the program *xmgrace* with the following command:

~~~
$ xmgrace-log x-nxy seq1.tkin
~~~

### 2.9. Coarse grain visualization to emphasize structural features

Kinetic folding plots (cf. subsection 2.8) usually produce a big amount of independent curves (500 in our example), one for each macro-state of the barrier tree. However, we optimized the RNA to exhibit specific structural features and thus want to visualize how often we observe this sub-structure in the ensemble of structures and the kinetic plots. Thus, we are collecting states that exhibit our structural features, i. e., ligand-binding stem or alternative stem, and summarize them into combined density curves. We implemented a *Perl* script called coarsify bmap.pl^1^ that performs this task. It can be applied to seq1.bar and seq1.tkin output as follows:

~~~
coarsify_regex.txt
# ?25(((((((((((……))))))))))) | ?26((((((((((……))))))))))
^.{25}\({11}\.{6}\){11}[\.\(\)]{11}|^.{26}\({10}\.{6}\){10}[\.\(\)]{11}
# ?2(((…((((((((…..)))))…)))…))) | ?2(((…((((((((…..))))…))))…)))”
^.{2}\({3}\.{3}\({8}\.{5}\){5}\.{3}\){3}\.{3}\){3}|^.{2}\({3}\.{3}\({8}\.{5}\){4}\.{3}\){4}\.{3}\){3}
$ perl coarsify_bmap.pl-regs coarsify_regex.txt-minh 30 \
-outdir coarse_30 seq1.bar seq1.tkin
~~~

This script merges macro-states of a given barrier tree in two ways: i) if the barrier height of a state is below the selected --minh value, it is merged to its neighbor and the population density of this neighbor is increased accordingly, and ii) if states contain similar structural elements, specified as regular expressions (coarsify_regex.txt), they are merged. Note that macro-states containing a different set of these structural elements are never merged although i) would be applicable. In the above example, all states are merged as --minh is larger than the energy band generated by *RNAsubopt*. However, the two specified regular expressions combine states that are compatible with the initial structural constraints of the design and keep the remaining landscape separate. The coarse-grained *barriers* and *treekin* output is written to the specified coarse 30/ subdirectory.

### 2.10. Kinetic simulation of an RNA with ligand interaction

Analyzing the influence of a ligand on the folding kinetics of a potentially binding-competent RNA molecule can, in its most general form, be a difficult problem. However, under certain conditions discussed below it is possible to use *treekin* (cf. Table 1) for this task. The effect of ligand addition can be simulated by declaring the binding-competent state *absorbing*, i. e., prevent any transitions out of it. This can be achieved by starting *treekin* with the population density of the last time point in seq1.tkin — i. e., the equilibrium distribution — and setting the-a option to the most stable binding-competent state:

~~~
$ grep-v “#” seq1.tkin | tail-n 1 | \
perl-ae ‘{for($i = 1; $i<scalar(@F); $i + +) {print” --p0 $i = $F[$i] “}}’ > t
$ treekin-m I `cat t`-f rates.out --t8 = 1E12-a 3 < seq1.bar > seq1_absorb.tkin
$ coarsify_bmap.pl-regs coarsify_regex.txt-minh 30 \-outdir coarse_30absorb seq1.bar seq1_absorb.tkin
$ rm t
~~~

First, the last time point in seq1.tkin is extracted and converted such that the output saved in t can be used as repeated --p0 parameter of *treekin*. Then, *treekin* is called and its output is stored in seq1_absorb.tkin, which is subsequently coarse grained. Finally, the temporary file t is removed. Visualization of the coarse-grained absorbing landscape is possible with the graph plotting tool *xmgrace* (cf. Table 1) by running:

~~~
$ xmgrace-log x-nxy coarse_30absorb/seq1_absorb.tkin
~~~

Using an absorbing state to model the ligand interaction is an approximation that is only reasonable under certain conditions. Irrespective of the properties of a specific ligand, a high ligand concentration as compared to the RNA concentration is assumed. In fact, absorbing states may be interpreted as an *infinite* ligand concentration leading to an immediate dimerization with the binding-competent RNA. Since ligands are usually much smaller molecules than their respective target RNA, this assumption is reasonable and ligand concentrations in the order of 1 mM are realistic in practice, though care has to be taken when dealing with toxic ligands like antibiotics.

The absorbing state assumption implies that the RNA – ligand complex has a rather low dissociation constant *K*_*d*_, or put differently, the dissociation rate coefficient *k*_off_ is low compared to its association rate coefficient *k*_on_. As pointed out by Wolfinger et al. in this special issue [20], in a working *(co-)transcriptional* RNA switch, the coefficients of the dimerization and the dissociation rate obey the criteria *k*_on_ *>* 1*/t*_apt_ and *k*_off_ ≪ 1*/t*_elong_, where *t*_apt_ is the duration during which the aptamer senses the ligand during transcription, and *t*_elong_ is the time required for the transcription of a single nucleotide. Under such conditions, the usage of an absorbing state to model the dimerization seems to be adequate. For *thermodynamic* switching behavior, *K*_*d*_ translates into a (negative) energy bonus *θ* = -*RT* ln *K*_*d*_ [21] awarded to all binding-competent structures. Let *E*_MFE_ be the MFE of the given sequence, and *E*_bc_ the free energy of the binding-competent state. Then, for the switch to work properly, *E*_bc_ + *θ* ≤ *E*_MFE_ ≤ *E*_bc_ must hold [22]. If the left-hand side of the inequality is significantly smaller, then the absorbing state model is a well-suited approach.

For many ligands of several different classes, aptamers with low *K*_*d*_ values in the order of 1 μM have been characterized, see [23] for a comprehensive summary. Though a precise *K*_*d*_ threshold cannot be given due to the concentration dependence, we hypothesize that such ligands behave in a way suited for this approximative approach.

## 3. Results

At the very beginning of a design process, necessary building blocks should be analyzed and evaluated to elucidate their properties. One of these blocks are RNA aptamers, as they cannot be generated by simply applying computational design methods. Here, we use an experimentally characterized RNA aptamer with known dissociation constant *K*_*d*_ and adaptive structural features, namely the well-known theophylline aptamer [17, 27, 28]. Alternatively, a novel aptamer could—at least in principle—be selected by performing an experimental protocol such as Systematic Evolution of Ligands by EXponential enrichment (SELEX) for the ligand of interest.

Next, a precise idea of how the desired RNA regulation mechanism should work is required. If it resembles a naturally occurring regulation mechanism, it is advisable to investigate its biological counterpart in detail before transferring the concept to a novel design. Figure 2 sketches the idea used to carry out the design step of this contribution. A given aptamer is extended in a way that an alternative structural conformation (ac) is formed in absence of the ligand. As the ligand is added, it reacts with the binding-competent conformation (bc) to form the ligand-bound conformation (lc), thereby stabilizing it and sequestering the alternative conformation (ac).

**Figure 2:**
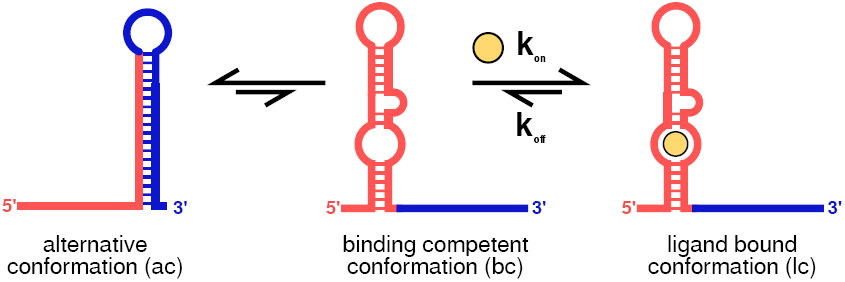
Graphical representation of the design idea. The system consists of two parts. In absence of the ligand, two conformations should dominate the structural ensemble. Depending on the design parameter, the alternative conformation (ac) should be higher populated than the binding competent (bc) one. Refolding rates between the two structural conformations depend on the energy barrier that separates them. Upon ligand addition, the bc gets trapped and the system should be shifted towards the ligand bound conformation (lc).

The mentioned downstream extension of the aptamer is necessary to introduce some degrees of freedom for the sequence sampler since the sequence of the aptamer itself is fixed. The insert needs to be long enough to sequester significant parts of the aptamer’s binding-competent structure. On the other hand, *short* inserts are preferable to avoid unforeseen interactions with the surrounding sequence context. Experiments by Ceres et al. [29] suggest that the ability of many aptamers to bind their respective ligand may be disrupted by solely opening their P1 (i. e., outermost) stem. However, we decided to introduce 22 nucleotides, corresponding to about half the length of the aptamer, to allow for an adequate thermodynamic stability and the complete sequestering of the aptamer structure by the alternative conformation.

We converted this model into a sequence and two structural constraints that represent ac and bc. If its structure had been resolved, the ligand-bound conformation could be taken into account as a third structural constraint. This would allow *RNAblueprint* to only generate sequences compatible to the structure of the dimer conformation. However, upon ligand binding, aptamers typically adapt complex tertiary interactions going beyond the scope of the classical secondary structure model. In case of theophylline, extensive stacking as well as the formation of base triples during ligand recognition have been observed [27, 28]. Such interactions cannot be handled by currently available secondary structure prediction and RNA design tools. A structural constraint modeling conformation lc is therefore omitted.

The functional model can be expressed as a combination of constraints such as structural requirements and various properties specifying the energy landscape, and the kinetic folding properties. An RNA sequence meeting these requirements as close as possible can be obtained by performing a local optimization approach. Such an approach includes i) the sampling of sequences with respect to a set of prescribed constraints, ii) the definition of a quality criterion through a proper objective function, and iii) an optimization method that decides whether to keep or reject a proposed solution.

We applied *RNAblueprint* to uniformly sample sequences that are compatible to the given sequence and structure constraints of the proposed design model, cf. subsection 2.1. The returned sequences need to be scored according to the design goal. Clearly, this design goal should include the evaluation of kinetic processes driving the implemented switching dynamics. However, these predictions are usually too demanding to be evaluated many times during optimization. Thus, we only use reasonable fast thermodynamic measures to ensure mandatory properties of the resulting kinetic processes [30].

On the thermodynamic level, we need to guarantee that the conformations of our model are exclusively present at least in the equilibrium. Given a sequence *x* and a compatible structure *φ*, one can calculate the corresponding Gibbs free energy *G*(*x* | *φ*) using the nearest neighbor model [31, 32]. The sequence – structure mapping is a one-to-many relation. Hence, one sequence can adapt a huge set of possible structures Φ called this sequence’s structure ensemble. In the equilibrium, the Boltzmann weight 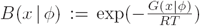 of a structure *φ* is proportional to its probability. Summing over all structures of the ensemble gives rise to the partition function *Z* = ∑_*φ*∊Φ_*B*(*x* | *φ*) of *x*. From that we can calculate the probability of *φ* with respect to the ensemble as

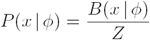

We utilize these properties to develop a novel objective function for the proposed model, cf. Figure 2. When adding the ligand to the system, we want to maximize the number of bound molecules, i. e., the probability of lc should ideally be one. As we do not have an explicit structural constraint of this state, we maximize the number of binding-competent structures in presence of the ligand, assuming the ligand is available in excess and immediately bound. In case of the theophylline aptamer, this precondition is fulfilled as its association rate constant *k*_on_ has been determined to be much higher than the dissociation rate constant *k*_off_ [28]. This is in accordance to its independently measured *K*_*d*_ of about 0.32 μM [17]. We therefore add an energy bonus of −9.22 kcal mol^−1^ to every secondary structure in the ensemble that contains the correctly folded — i. e., binding-competent — theophylline aptamer.

By maximizing the probability of bc in the presence of the ligand, we favor the conversion to the ligand-bound conformation lc. In contrast, ac should be highly populated in absence of the ligand. However, no ligand binding is possible if the RNA molecule exclusively adapts ac as only bc induces a high binding affinity of the ligand for the RNA molecule. It is therefore necessary to establish a balance between ac and bc where bc must always be present. We combined all these assumptions into the novel objective function

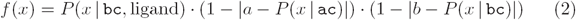

where *a, b* ∊ (0, 1), *a* + *b* ≤ 1 are the target probabilities of the alternative conformation and binding-competent conformation, respectively. This function is maximized as all terms tend to one. We set *a* = 0.7 and *b* = 0.3 for the discussed example. We describe the details on how to calculate the individual terms of the objective function given above utilizing the constraint framework of the *ViennaRNA* package in subsection 2.3 and subsection 2.4.

To perform a local optimization procedure searching for sequences optimal with respect to the derived objective function (2), we chose to harness a scripting library called *RNAsketch*, which is available as interface to the sequence sampler *RNAblueprint* (cf. Table 1). *RNAsketch* offers ready-to-use implementations of several well-known optimization strategies. To tackle the presented design problem, we implemented a *Python* script that performs adaptive walks with randomly chosen steps of varying size — ranging from point mutations to full resampling of the sequence — until the score evaluated with the designed objective function (2) stays minimal. This approach has been found to converge relatively fast towards reasonable results for other objectives [10]. Our implementation^2^ and the corresponding commands including the inputs are available online to serve as an example of use for *RNAsketch*.

The described local optimization procedure is capable of producing many potential solutions in a relatively short amount of time. As the returned scores contain no additional information but the three probabilities, we developed an *in silico* analysis pipeline to visualize additional properties of the obtained sequences, facilitating a consecutive ranking and filtering step. First and foremost, we need to verify the kinetic properties of our obtained solutions, a usually very expensive and time-consuming task. In the following, we discuss this process for an example sequence^3^.

For a more complete picture of the energy landscape, we need to investigate the structural states our example sequence will likely fold into. *RNAsubopt* is applied to generate all suboptimal structures up to 22.6 kcal mol^−1^ above the sequence’s minimum free energy, cf. subsection 2.5. The number of possible structures grows exponentially with sequence length and is approximately 225 million for the chosen energy range of 22.6 kcal mol^−1^, resulting in a 16 GB large file. To reduce the number of generated suboptimal structures, and thereby speed up all subsequent steps, it is possible to skip all structures containing lonely pairs, i. e., helices of length one, generating only so-called canonical structures. This reduces the number of states to approximately 6.7 million, and the file size to 459 MB. However, for the shown example, this also excludes the predicted MFE structure, cf. most populated structures in the equilibrium in Figure 3. The previous ground state containing the alternative structural element is only the fourth-stable state while the MFE structure contains the binding-competent aptamer. Of course, this has a dramatic impact on the simulated kinetics, cf. Figure 3.

**Figure 3:**
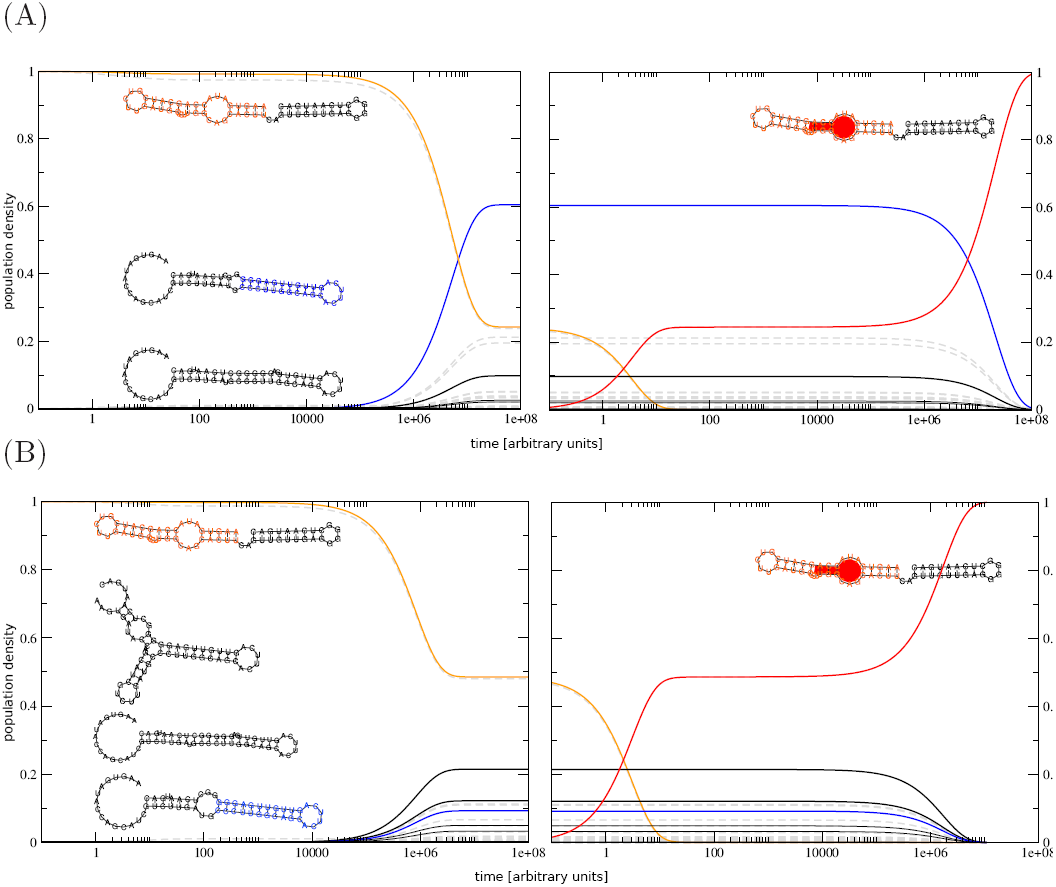
Simulated kinetics using (A) the complete and (B) a reduced structure ensemble by avoiding lonely pairs. In both cases, the simulation is started with the complete population in a structural state that contains the binding competent aptamer structure (orange). The left part of each plot shows the dynamics until the system is equilibrated, whereas the right part depicts the simulated systems kinetics after ligand addition. Dashed gray lines indicate the system’s kinetics without coarse graining. By design, the population density in the equilibrium of all structures containing the alternative (blue) and the binding-competent (orange) structural element should be 0.7 and 0.3, respectively. Colored lines display the coarse-grained kinetics where states containing specific structural elements are merged. For the most prominent states, the secondary structure corresponding to their respective stable representative are shown using the same color. Despite the similarities of the representatives of the blue and the thick black curves, they were not merged because the alternative structural element was required to have a perfectly stacked stem of length at least 11 nt.

To assess the impact of the “no lonely pairs” (--noLP) heuristics more profoundly, the procedure described in subsection 2.6 has been applied to the example sequence. By enumerating all structures up to 10 kcal mol^−1^ above the MFE, one obtains a structure coverage of 99.9%. Here, the identified fraction of canonical structures is only 43%, so the vast majority of structures that would likely be encountered in the simulation are removed when applying the --noLP heuristics. It is clear that in this case, the reduction to the canonical structure ensemble leads to a strong bias.

In contrast, other sequences have much higher fractions of canonical structures. The example sequence^4^ has the same length (64 nt) and GC content (51%) as the previous one, but exhibits a predominantly canonical ensemble (96% of the structures).

In any case, further coarse graining of the structure landscape is mandatory. We apply the program *barriers* which implements a flooding algorithm and abstracts the structure landscape to a selected number of macro-states, each represented by a local minimum of the landscape (subsection 2.7). Transition rates from each of these macro-states to all other ones are then estimated and subsequently used to predict the folding kinetics.

It is possible that multiple macro-states exhibit structural features such as the structure of bc or the stem of ac. Thus, for better visualization we merged states that exhibit certain structural features by implementing *coarsify bmap.pl*, cf. subsection 2.9. Based on the resulting landscape and the processed transition rates, *treekin* has been invoked to simulate the single-molecule folding kinetics, cf. subsection 2.8. A visualization of the output shows the expected population density of the two designed structural states ac and bc after the equilibrium has been reached, cf. Figure 3A. This way, we verified that the estimate based on partition function folds — as used during the optimization process — matches the results of the kinetic simulation even in absence of the ligand and when the full *RNAsubopt* output, including non-canonical structures, is used.

When sketching the design (cf. Figure 2), we assumed that the RNA – ligand complex has a rather low dissociation rate coefficient *k*_off_ compared to its association rate coefficient *k*_on_. For the theophylline aptamer, this is in accordance with published rates of (0.07 ± 0.02) sec^−1^ and (1.7 ± 0.2) × 10^5^ M^−1^ sec^−1^ at 25^°^C for *k*_off_ and *k*_on_, respectively [28]. We therefore modeled the effect of ligand addition by starting *treekin* with the population density of the equilibrium and making the binding-competent state absorbing, cf. subsection 2.10. Visualization of the coarse-grained absorbing landscape shows that after about 9 × 10^6^ AU, which can be mapped to approximately 45 sec [33], 50% of the RNA molecules are in the ligand-bound state, cf. Figure 3.

## 4. Discussion

During the development of our software pipeline we realized that, until recently, there mainly existed two kinds of publications. One created by wet lab researchers, focused on an experimental testing setup as well as functional and analytical tests. They frequently missed the possible advances of *in silico* tools and their valuable predictive power. In contrast, publications written by researchers mainly working on computational biology often comprised sophisticated biophysical methods, great computational details, and a huge variety of mathematical and algorithmic tricks, but were frequently neglecting the aspect of biological applicability.

A main reason for this situation is that most of the RNA design programs available use predefined terms in the objective function as well as a fixed optimization procedure [22], and thus are inflexible and not customizable enough to be applied and adopted to the huge amount of considerably varying design scenarios. The *RNAblueprint* approach[10] decouples sampling of sequences compatible to one or more structural constraints from the subsequent optimization procedure, which gives the user the full flexibility to implement novel and innovative objectives.

Computational design studies are often missing the bigger context, such as the initial analysis of the system, suggestions for experimental testing or the design of proper controls. However, experimental validation is not a straightforward task and needs to be carefully planned already during the design process. This includes extensive *in vitro* or *in vivo* studies, or preferable both. To really gain knowledge about the device’s mechanism and about potential mistakes or pitfalls in case of dysfunctionality, a purely qualitative answer will not be sufficient. Therefore, a complete testing pipeline should include the determination of structures, binding affinities, or elucidate kinetic properties. Smartly designed positive and negative controls are also helpful to reveal important properties of the newly generated RNA device. Ideally, these controls will unveil quantitative answers about the mechanistic details, the actual structures of the RNA or even about kinetic aspects like co-transcriptional dependency or ligand affinity.

In this contribution, we described in detail the *de novo* design of a ligand-sensing riboswitch that adapts two alternative conformations. Depending on the presence of the ligand, either a binding competent state, or a specified alternative structural conformation is dominating the ensemble. This riboswitch design can easily be extended, e. g., to perform regulatory tasks in a host cell such as translational or transcriptional regulation of a downstream target gene. A translational riboswitch for instance will probably contain a Ribosome Binding Site (RBS) which is sequestered in the inactive state. This can be included easily by specifying the appropriate sequence constraints and further objectives such as the accessibility of the RBS in both conformations.

For such a purpose, it is important to distinguish between two types of switching behaviors One type of riboswitches which are capable to switch on and off during the entire lifetime of the molecule. The other switches are fixed after a certain time of sensing disregarding future changes in the ligand concentration. For the latter, switching is only possible through RNA decay and repeated transcription.

If switching is possible at any time, fast response times to ligand changes are obtained. However, the individual states of the molecules remain fuzzy as not all of them will adapt the desired structure, leading to the observation of background activity. In our example, we provoked such a behavior by targeting a 70:30 ratio of alternative to binding-competent conformation, and indeed observed a quick refolding process upon addition or removal of the ligand.

Alternatively, we could generate a “one way switch” where the state decision is only possible during a specific window. Thereafter, the chosen state is stabilized, either by a kinetic folding trap or by ongoing molecular processes such as translation. Once decided, individual molecules cannot revert their choice within reasonable time, even if the ligand dissociates or is removed from the system [34]. Therefore, it must be ensured that the competing states are populated during the decision window. This method has the advantage of obtaining distinct states with very little noise. However, the response times to ligand changes are quite long as they depend on RNA decay and the transcription speed.

At a first glance, the design model we proposed in Figure 2 seems to be rather easy. However, it is not straight forward to develop an experimental setup that is able to determine if the target ratio of 70:30 of the two conformations is reached in the equilibrium or not. Sophisticated approaches such as single-molecule FRET and NMR have been applied to determine the structure and energy landscape of natural riboswitches[35, 36]. Both referenced studies revealed that more than the presumably two dominating states are adapted depending on environmental conditions, i. e., Mg^2+^ concentration and temperature. This might be the case for our designs as well although we optimized them towards two alternative states only. Furthermore, it is important to note that the presented *in silico* results are estimates. For instance, the target ratio of 70:30 might be achieved perfectly by the optimization procedure and predicted by kinetics simulations. However, the results are extremely sensitive to the underlying energy parameters. Those are measured under specific experimental conditions for rather short structural elements. For more complex structures, i. e., those containing large or even multi loops, estimates are utilized to deter-mine a structure’s energy [31]. If the experimental conditions used to determine the energy parameters and those used in the *in vitro* or *in vivo* testing environment vary significantly, discrepancies of prediction and measurement are an inevitable effect.

Depending on the research question, neglecting structures with lonely pairs can give valuable insights into the studied system while dramatically speeding up the prediction process. The reason the analysis fails for the exemplary sequence presented in this work is that many of its low-energy structures contain lonely pairs and are therefore excluded when enabling the heuristics. The effect is dramatic here as even the MFE structure is not canonical.

A method to assess the importance of lonely pairs for the simulations has been developed and shown to correctly predict the consequences of noLP heuristics. In general, it is advisable to always consider the fraction of canonical base pairs before resorting to the heuristic. Another advantage of conducting this additional analysis is the proper estimation of the energy band width required to achieve a high coverage of the structure ensemble during the enumeration. This information is useful to improve both, the performance of the simulation as well as the quality of the results, even when not utilizing the noLP heuristics. Nevertheless, re-running the analysis of promising candidates with a full structure ensemble is advisable to assure correct results if the required resources are available.

Many of the techniques used in this work implicitly make simplifying assumptions about the processes involved in RNA switching. For example, the soft constraint framework is a considerable abstraction of the binding process in at least two ways. Firstly, it models a binary binding behavior in that the ligand either perfectly fits an RNA structure and gets the full binding energy bonus, or it does not bind to the structure at all. In reality, small variations in the binding domain may lead to an altered binding energy instead. Secondly, any structure exhibiting the binding site receives the full stabilizing energy contribution, neglecting the effect of the ligand concentration and assuming infinite reaction rates. During the kinetics simulation, a similar behavior is achieved by declaring the binding-competent macro-state absorbing, i. e., the dissociation of the ligand is not possible at all.

While these may be adequate assumptions for ligands with a high binding affinity present in an excessive concentration, it may lead to over-estimation of the fraction of RNA – ligand complexes in situations where the association rate becomes the bottleneck of the dimerization reaction. In such cases, one should resort to more sophisticated models considering these rates as well as the ligand’s concentration [37]. An efficient implementation of this approach that can readily be applied to ligand-aware co-transcriptional folding is published in this special issue [20].

When analyzing our designed switch *in silico*, we started the kinetic simulation of the ligand-free environment with all molecules in the binding – competent state. Thereby, we ensured that even in this worst scenario possible, the system quickly recovers to the defined ratio of alternative state and binding-competent state. To obtain better estimates of the switching times, starting with various other distributions depending on the application might be preferable.

In case of an *in vitro* experiment, the protocol would probably envisage to first heat up the solution to completely untangle the RNA structures and then quickly put the solution on ice until the ligand is added. A similar cooling experiment could be performed *in silico* by performing Boltzmann-weighted structure sampling from an ensemble at high temperature and using the resulting distribution of states as starting point for a subsequent kinetic simulation. In contrast, when using the generated riboswitch *in vivo*, it is likely co-transcriptionally folded within the cell. Therefore, it is advisable to obtain the initial distribution by applying a co-transcriptional folding approach which simulates the RNA’s elongation process until the binding-competent part of the structure is fully transcribed. A software capable of this type of analysis is, for example, *BarMap* [38].

In this contribution, we aimed to generate a general ligand-triggered riboswitch which can be extended to control regulation mechanisms, such as transcriptional or translational control of a downstream target gene. We devised a functional model of such a riboswitch and successfully implemented a design approach to *de novo* generate RNA sequences that fulfill the prescribed properties. The proposed pipeline consists of several modular pieces which can be easily adopted or exchanged in case of varying needs. This includes the flexible sequence sampling engine *RNAblueprint*, a novel objective function to thermodynamically describe important features of the mechanism, the optimization approach and, finally, the *in silico* analysis pipeline to verify kinetic properties of the system.

## 5. Acknowledgments

This work has been supported by the European Commission under the Environment Theme of the 7th Framework Program for Research and Technological Development (Grant agreement number 323987), the Austrian science fund FWF project F43 “RNA regulation of the transcriptome”, the German Network for Bioinformatics Infrastructure (de.NBI) by the German Federal Ministry of Education and Research (BMBF; support code 031A538B), and by the German Research Foundation (DFG; grant STA 850/15-2)

We thank Christina Wagner for fruitful discussion, Manuela Geiß for assistance with mathematical issues and our private experimental testing help desk Mario Mörl.

## Abbreviations

UTR: untranslated region
mRNA: messenger RNA
SELEX: Systematic Evolution of Ligands by EXponential enrichment
MFE: minimum free energy
RBS: Ribosome Binding Site

## Footnotes

1 https://github.com/ViennaRNA/BarMap/

2 design-ligandswitch.py

3 AAGUGAUACCAGCAUCGUCUUGAUGCCCUUGGCAGCACUUCAGUUGUUGAGGGGGCUCAAUGAC

4 GUAAGAGAGGCCGCGCACAACUUUCCUACUGUUCGAAAGGUAGGAGCGCUGUCAACUUACAUGG

## References

[1] R. R. Breaker, Prospects for Riboswitch Discovery and Analysis, Molecular Cell 43 (6) (2011) 867–879. doi:10.1016/j.molcel.2011.08.024.

[2] C. C. Fowler, E. D. Brown, Y. Li, A FACS-based approach to engineering artificial riboswitches., Chembiochem 9 (12) (2008) 1906–1911. doi:10.1002/cbic.200700713.

[3] J. E. Weigand, M. Sanchez, E.-B. Gunnesch, S. Zeiher, R. Schroeder, B. Suess, Screening for engineered neomycin riboswitches that control translation initiation, RNA 14 (1) (2008) 89–97. doi:10.1261/rna.772408.

[4] J. E. Weigand, B. Suess, Tetracycline aptamer-controlled regulation of pre-mRNA splicing in yeast, Nucleic Acids Research 35 (12) (2007) 4179–4185. doi:10.1093/nar/gkm425.

[5] C. Schneider, B. Suess, Identification of RNA aptamers with riboswitching properties, Methods 97 (2016) 44–50. doi:10.1016/j.ymeth.2015.12.001.

[6] A. A. Green, P. A. Silver, J. J. Collins, P. Yin, Toehold Switches: De-Novo-Designed Regulators of Gene Expression, Cell 159 (4) (2014) 925–939. doi:10.1016/j.cell.2014.10.002.

[7] M. Etzel, M. Mörl, Synthetic Riboswitches: From Plug and Pray toward Plug and Play, Biochemistry 56 (9) (2017) 1181–1198. doi:10.1021/acs.biochem.6b01218.

[8] A. Churkin, M. D. Retwitzer, V. Reinharz, Y. Ponty, J. Waldispühl, D. Barash, Design of RNAs: Comparing programs for inverse RNA folding, Briefings in Bioinformatics (2017) bbw120 doi:10.1093/bib/bbw120.

[9] F. J. Isaacs, D. J. Dwyer, C. Ding, D. D. Pervouchine, C. R. Cantor, J. J. Collins, Engineered riboregulators enable post-transcriptional control of gene expression, Nature Biotechnology 22 (7) (2004) 841–847. doi:10.1038/nbt986.

[10] S. Hammer, B. Tschiatschek, C. Flamm, I. L. Hofacker, S. Findeiß, RN-Ablueprint: flexible multiple target nucleic acid sequence design, Bioinformatics 33 (18) (2017) 2850–2858. doi:10.1093/bioinformatics/btx263.

[11] N. Metropolis, A. W. Rosenbluth, M. N. Rosenbluth, A. H. Teller, E. Teller, Equation of State Calculations by Fast Computing Machines, The Journal of Chemical Physics 21 (6) (1953) 1087–1092. doi:10.1063/1.1699114.

[12] W. K. Hastings, Monte Carlo sampling methods using Markov chains and their applications, Biometrika 57 (1) (1970) 97–109. doi:10.1093/biomet/57.1.97.

[13] A. Taneda, Multi-objective optimization for RNA design with multiple target secondary structures, BMC Bioinformatics 16 (1) (2015) p280. doi:10.1186/s12859-015-0706-x.

[14] I. L. Hofacker, W. Fontana, P. F. Stadler, L. S. Bonhoeffer, M. Tacker, P. Schuster, Fast folding and comparison of RNA secondary structures, Monatshefte fu¨r Chemie / Chemical Monthly 125 (2) (1994) 167–188. doi:10.1007/BF00818163.

[15] C. Höner zu Siederdissen, S. Hammer, I. Abfalter, I. L. Hofacker, C. Flamm, P. F. Stadler, Computational design of RNAs with complex energy landscapes, Biopolymers 99 (12) (2013) 1124–1136. doi:10.1002/bip.22337.

[16] R. Lorenz, I. L. Hofacker, P. F. Stadler, RNA folding with hard and soft constraints, Algorithms for Molecular Biology 11 (8) (2016). doi:10.1186/s13015-016-0070-z.

[17] R. D. Jenison, S. C. Gill, A. Pardi, B. Polisky, High-resolution molecular discrimination by RNA, Science 263 (5152) (1994) 1425–1429.

[18] Y. Ding, C. Y. Chan, C. E. Lawrence, RNA secondary structure prediction by centroids in a boltzmann weighted ensemble, RNA 11 (8) (2005) 1157–1166.

[19] C. Flamm, I. Hofacker, Beyond energy minimization: approaches to the kinetic folding of RNA, Chemical Monthly 139 (2008) 447–457.

[20] M. T. Wolfinger, C. Flamm, I. L. Hofacker, Efficient computation of co-transcriptional RNA-ligand interaction dynamics, Methods Submitted.

[21] M. Wachsmuth, S. Findeiß, N. Weissheimer, P. F. Stadler, M. Mörl, De novo design of a synthetic riboswitch that regulates transcription termination, Nucleic Acids Research 41 (4) (2013) 2541–2551. doi:10.1093/nar/gks1330.

[22] S. Findeiß, M. Wachsmuth, M. M¨orl, P. F. Stadler, Design of transcription regulating riboswitches, Methods Enzymol 550 (2015) 1–22. doi:10.1016/bs.mie.2014.10.029.

[23] S. Klussmann (Ed.), The Aptamer Handbook: Functional Oligonucleotides and Their Applications, Wiley-VCH, 2006. doi:10.1002/3527608192.

[24] C. Flamm, I. L. Hofacker, P. F. Stadler, M. T. Wolfinger, Barrier trees of degenerate landscapes, Zeitschrift für Physikalische Chemie 216 (2/2002) (2002). doi:10.1524/zpch.2002.216.2.155.

[25] M. T. Wolfinger, W. A. Svrcek-Seiler, C. Flamm, I. L. Hofacker, P. F. Stadler, Efficient computation of RNA folding dynamics, Journal of Physics A: Mathematical and General 37 (17) (2004) 4731–4741. doi:10.1088/0305-4470/37/17/005.

[26] R. Lorenz, S. H. Bernhart, C. H. z. Siederdissen, H. Tafer, C. Flamm, P. F. Stadler, I. L. Hofacker, ViennaRNA Package 2.0, Algorithms for Molecular Biology 6 (1) (2011) p26. doi:10.1186/1748-7188-6-26.

[27] G. R. Zimmermann, R. D. Jenison, C. L. Wick, J.-P. Simorre, A. Pardi, Interlocking structural motifs mediate molecular discrimination by a theophylline-binding RNA, Nature Structural Biology 4 (8) (1997) 644–649. doi:10.1038/nsb0897-644.

[28] F. M. Jucker, R. M. Phillips, S. A. McCallum, A. Pardi, Role of a heterogeneous free state in the formation of a specific RNA-theophylline complex., Biochemistry 42 (9) (2003) 2560–2567. doi:10.1021/bi027103+.

[29] P. Ceres, J. J. Trausch, R. T. Batey, Engineering modular ON RNA switches using biological components, Nucleic Acids Research 41 (22) (2013) 10449–10461. doi:10.1093/nar/gkt787.

[30] S. Badelt, S. Hammer, C. Flamm, I. L. Hofacker, Chapter Eight - Thermodynamic and Kinetic Folding of Riboswitches, in: S.-J. C. Burke-Aguero, D. H. (Eds.), Methods in Enzymology, Vol. 553 of Computational Methods for Understanding Riboswitches, Academic Press, 2015, pp. 193–213.

[31] D. H. Mathews, M. D. Disney, J. L. Childs, S. J. Schroeder, M. Zuker, D. H. Turner, Incorporating chemical modification constraints into a dynamic programming algorithm for prediction of RNA secondary structure., Proc Natl Acad Sci U S A 101 (19) (2004) 7287–7292. doi:10.1073/pnas.0401799101.

[32] D. H. Turner, D. H. Mathews, NNDB: the nearest neighbor parameter database for predicting stability of nucleic acid secondary structure., Nucleic Acids Res 38 (Database issue) (2010) D280–D282. doi:10.1093/nar/gkp892.

[33] B. Sauerwine, M. Widom, Folding kinetics of riboswitch transcriptional terminators and sequesterers, Entropy 15 (8) (2013) 3088–3099. doi:10.3390/e15083088.

[34] G. Quarta, K. Sin, T. Schlick, Dynamic energy landscapes of riboswitches help interpret conformational rearrangements and function., PLoS Comput Biol 8 (2) (2012) pe1002368. doi:10.1371/journal.pcbi.1002368.

[35] A. Reining, S. Nozinovic, K. Schlepckow, F. Buhr, B. Fürtig, H. Schwalbe, Three-state mechanism couples ligand and temperature sensing in riboswitches, Nature 499 (7458) (2013) 355–359. doi:10.1038/nature12378.

[36] C. Manz, A. Y. Kobitski, A. Samanta, B. G. Keller, A. Jschke, G. U. Nienhaus, Single-molecule FRET reveals the energy landscape of the full-length SAM-i riboswitch, Nature Chemical Biology 13 (11) (2017) 1172–1178. doi:10.1038/nchembio.2476.

[37] F. Kühnl, P. F. Stadler, S. Will, Tractable RNA–ligand interaction kinetics, BMC Bioinformatics 18 (S12) (2017). doi:10.1186/s12859-017-1823-5.

[38] I. L. Hofacker, C. Flamm, C. Heine, M. T. Wolfinger, G. Scheuermann, P. F. Stadler, BarMap: RNA folding on dynamic energy landscapes, RNA 16 (2010) 1308–1316.

